# Mental geometry of perceiving 3D size in pictures

**DOI:** 10.1101/2020.04.04.025759

**Authors:** Akihito Maruya, Qasim Zaidi

## Abstract

We show that the classical problem of 3D size perception in obliquely viewed pictures can be understood by comparing human performance to the optimal geometric solution. A photograph seen from the camera position, forms the same retinal projection as the physical 3D scene, but retinal projections of sizes and shapes are distorted in oblique viewing. For real scenes, we previously showed that size and shape inconstancy result despite observers using the correct geometric back-transform, because some retinal images evoke misestimates of viewing elevation. Now, we examine how observers estimate 3D sizes in oblique views of pictures of objects in different poses on the ground. Compared to the camera position estimates, sizes in oblique views were seriously underestimated for objects at fronto-parallel poses, but there was almost no change for objects perceived as pointing towards the viewer. The inverse of the function relating projected length to pose, camera elevation and viewing azimuth, gives the optimal correction factor for inferring correct 3D lengths if the elevation and azimuth are estimated accurately. Empirical correction functions had similar shapes to optimal, but lower amplitude. Measurements revealed that observers systematically underestimated viewing azimuth, similar to the fronto-parallel bias for object pose perception. A model that adds underestimation of viewing azimuth to the geometrical back-transform, provided good fits to estimated 3D lengths from oblique views. These results add to accumulating evidence that observers use internalized projective geometry to perceive sizes, shapes and poses in 3D scenes and their pictures.

## Introduction

A major part of scene understanding for locomotion, hunting, gathering, shelter, and even aesthetic appreciation, consists of judging poses, sizes and shapes of objects. Since projective geometry determines retinal and camera images of objects, a brain or machine could calculate accurate poses and sizes if it could invert the projection. The inversion back from 2D to 3D can generate unique estimates of poses and sizes of a known object if viewing parameters are ascertained independently. Building on previous work (Koch, Baig & Zaidi, 2018; Maruya & Zaidi, JOV In press), we tackle the classical problem of the perception of object sizes (and shape aspect ratios) from different views of pictures of 3D scenes (Boring, 1964; Gombrich, 1972; Perkins, 1973; Hagen, 1974; Hagen, 1976; Mulholland et al, 1980; Wallach et al, 1986; Cutting, 1987; Niall et al, 1990; Yang et al, 1999; Vishwanath et al, 2005; Todorovorić, 2008), and identify the geometric operations that are used for estimation. The results help to understand not just picture perception, but also the mental geometry that is essential for understanding 3D scenes.

Koch, Baig & Zaidi (2018) showed that human observers are very accurate in judging 3D poses of objects on the ground, albeit with a fronto-parallel bias, and their results can be explained by a model using the back-transform of the projective geometry functions that maps 3D poses to retinal orientations, with just one free parameter for the bias. There was exceedingly close correspondence across observers (Pair-wise correlations mean = 0.9934, sd = 0.0037), and individual results all fit the same geometric back-transform function, suggesting strongly that object-pose estimation, which has been a critical function for millions of years of evolution, could be based on internalized geometric knowledge. Pictorial representation, on the other hand, is less than forty thousand years old, so we tested the possibility that observers estimate 3D poses in photographs by using the same geometrical operations as for real scenes, despite retinal images being further distorted by projection to oblique viewpoints. The model based on back-projections from retinal orientations, predicts rigid rotation of perceived poses, so that objects pointing at an observer in the real scene should also be seen as pointing at the observer in obliquely viewed photographs of the scene. Pose estimates corresponded almost perfectly (R^2^>0.99) with the rotation prediction, thus providing support for the use of these geometrical operations. These results also confirmed that the “pointing at you” pictorial phenomena can be explained simply in terms of back-projections from vertical retinal orientations without invoking any previously postulated pictorial spaces or operations (Kennedy, 1974; Ward, 1976; Pierroutsakos et al, 1998; Yang et al, 1999; Heyer, 2003; Vishwanath et al, 2005; Koenderink et al, 2004 and 2011; Pagel, 2017). This explanation fits with Cézanne’s observation in his letter to Émile Bernard dated 15 April 1904: “Lines parallel to the horizon give breadth, a section of nature, or if you prefer, of the spectacle spread before our eyes Lines perpendicular to that horizon give depth.”

Using the same research strategy, Maruya & Zaidi (JOV In press) examined 3D size perception in real scenes as a function of the pose of the object. Unlike the usual experimental method of varying distance from the observer to study size constancy (Brunswik, E., 1944; Gilinsky, 1951 and 1955; Carlson, V. R., 1960; Norman et al, 1996; Beusmans, 1998; Ross, H. E., & Plug, C., 1998; Loomis et al, 1992, 1999, and 2002), experimentally varying object pose changes the shape projected on the retina, so requires quite different compensations for projective distortions. Observers compensated for projective shortening as a function of 3D pose, but not sufficiently for objects pointing towards or away from the observer, which are the poses that project to the shortest retinal sizes. Modeling the empirical correction as a function of the optimal correction, revealed that perceived sizes in 3D scenes are inconstant despite observers using the correct geometric back-transform, because the retinal image evokes a slant illusion that reduces the compensation.

The illusory rotation caused by oblique viewing of pictures (Koch, Baig, Zaidi, 2018), makes poses invariant to viewing azimuth, but does lead to perceived variation in sizes and shapes, especially aspect ratios. Therefore, in this study, we test whether 3D size estimates in oblique views of pictures still follow from the geometric back-transform, and if there are simple factors that can explain estimation errors. Whereas projection from a real scene to the retina reduces sizes most along the axis pointing to the observer, viewing the picture obliquely (or equivalently tilting the picture around the vertical axis) reduces sizes most along the fronto-parallel axis. So we were particularly interested in any role played by the perceived tilt of the picture with respect to the observer, since that had no effect on 3D pose estimation from pictures, which used only retinal orientations. We found that 3D sizes at fronto-parallel poses are seriously underestimated in oblique views compared to the frontal view. By contrast, there was almost no change for objects perceived as pointing to or from the viewer. We were able to model observers’ corrections for size as a function of pose, by using the optimal geometric back-transform with a multiplicative parameter for the systematic underestimation of the tilt of the display. The underestimation was confirmed by perceived tilt measurements, and is similar to the fronto-parallel bias for object pose perception. The excellent fit of the model shows that all observers use the correct back-transform from projective geometry, indicating the use of ingrained geometrical operations just as in pose estimation, but the perceived tilt of the picture plays a role in size estimation unlike in pose estimation. Since aspect ratios of 3D shapes depend on relative sizes along different axes, those aspects are subject to the same picture tilt distortions as perceived 3D sizes.

## Size Estimates of 3D objects at Oblique Views of pictures

### Methods

Using Blender, we created a blue rectangular 3D parallelepiped (test stick) lying on the center of a dark ground, and a yellow vertical 3D cylinder (measuring stick) standing on the test stick. Blue parallelepipeds were presented in one of 16 poses from 0° to 360° every 22.5°, of which poses in one quadrant are shown in Figure 1a. The line of sight through the center of the ground was designated the 90°-270° axis, and the line orthogonal to it as the 0°-180° axis. Parallelepiped were 10, 8 or 6 cm long with a 3×3 cm cross-section. Images were displayed on a 22-inch DELL SP2309W Display. Matlab and Psychtoolbox were used to display the stimuli, run the experiments, and analyze the data. The observer’s viewing position was fixed by using a chinrest so that the center of the monitor was viewed with an elevation angle of 15° at a distance of 1.0m, matching the rendering parameters of the Camera in Blender. Displayed sizes in the Blender rendered images were calibrated against exact geometrical derivations to ensure accuracy of the simulations (see Maruya & Zaidi, In press). Measurements were made from five viewpoints, with the screen in frontal (0°) position, and tilted at azimuths of ±30° and ±60°around a vertical axis. Derived lengths of retinal projections of the parallelepipeds are shown as a function of pose for fronto-parallel and oblique views of the display screen in Figure 1b. The derivation is detailed in the Appendix, and illustrated in Figure A1. For a parallelepiped of length (*L*_3*D*_), the projected length on the retina (*L_r_*) changes with pose (Ω) as a distorted sinusoid affected by the values for viewing elevation=*ϕ_c_*, viewing azimuth (equivalent to display tilt) =*ϕ_v_*, focal length of camera=*f_c_*, focal length of eye = *f_c_*, distance from the object=*d_c_*, and distance from the center of the display=*d_v_*:

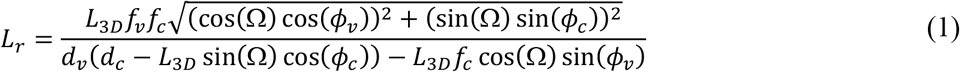

**Figure 1.**
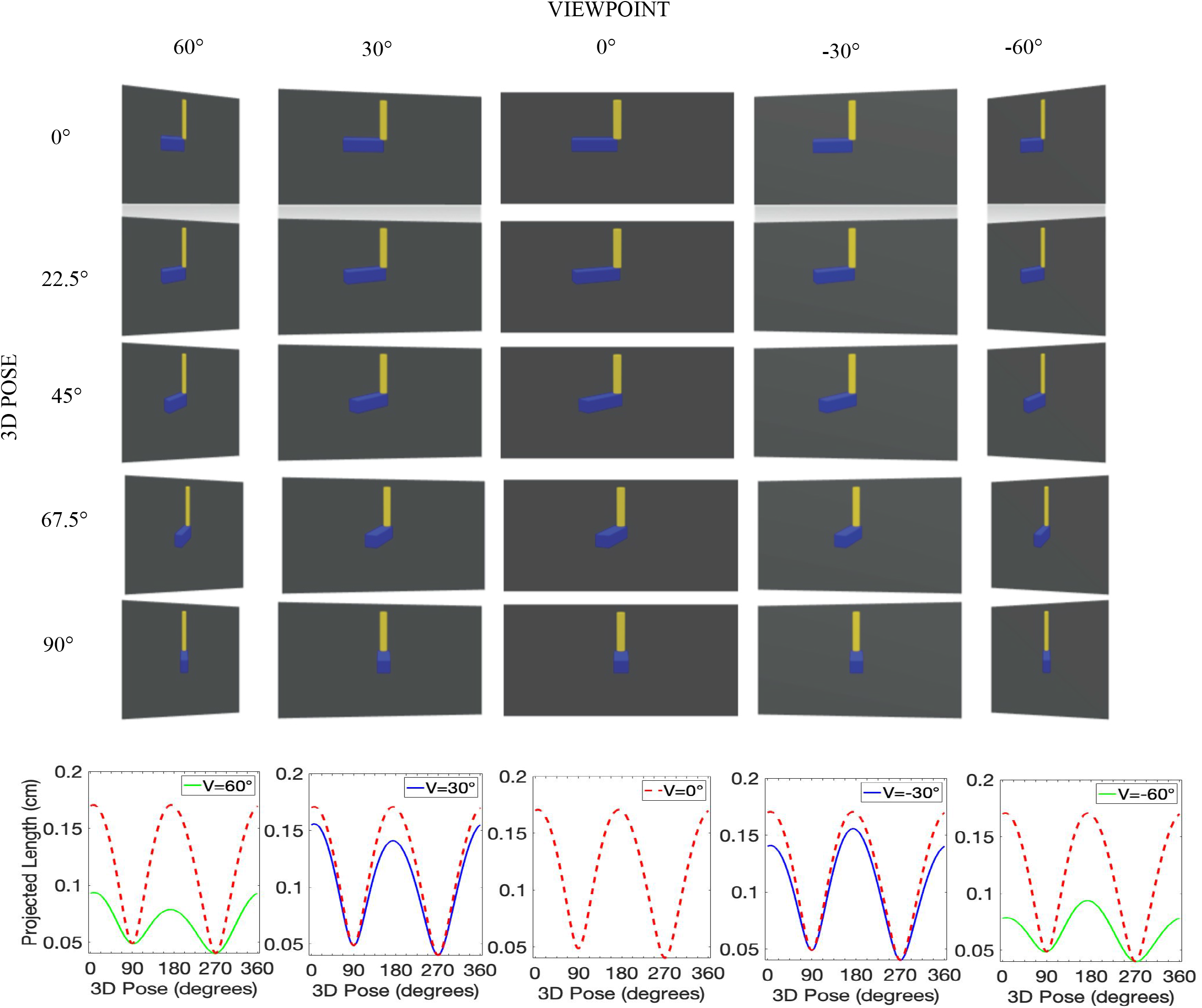
Projected images and lengths of Experiment 1 stimuli. (a) Blue rectangular 3D parallelepiped (test stick) of fixed length lying on the center of a dark ground, and a yellow vertical 3D cylinder (measuring stick) of adjustable length standing on the test stick. Blue parallelepipeds were presented in one of 16 poses from 0° to 360° every 22.5°, of which poses in one quadrant are shown in Figure 1(a). The line of sight through the center of the ground is designated the 90°-270° axis, and the line orthogonal to it as the 0°-180° axis. Parallelepiped lengths were 10, 8 or 6 cm with a 3×3 cm cross-section. Images were displayed on a 22-inch display. The monitor was viewed with an elevation angle of 15° at a distance of 1.0m, from five viewpoints: frontal (0°), and tilted at azimuths of ±30° and ±60°around a vertical axis. Observe that projected length is shortest for the 90° pose in the 0° viewpoint but it changes little across viewpoints, whereas the length for the 0° pose varies most across viewpoints. (b) Derived lengths of retinal projections of the parallelepipeds are shown as a function of pose for the five views of the display screen. The derivation is detailed in the Appendix, and illustrated in Figure A1.

However, the projected length *L_mr_* on the retina of the vertically oriented cylinder of physical length *L*_3*Dm*_, stays invariant with object pose and display tilt because the 3D object and the picture plane are both rotated around the vertical axis of the cylinder:

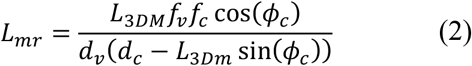

Observers were instructed to equate the physical lengths of the two limbs by pushing buttons to adjust the height of the measuring stick between 2.75 and 12 cm. There were no time limits. Randomly ordered trials across all three sizes and all sixteen poses were run in blocks for each of the five tilts of the display, and repeated in three sets. Observers were allowed to take a break between sets.

6 observers with normal or corrected vision participated. Viewing was binocular because it is the natural condition for looking at pictures, and Koch et al (2018) had not found any difference between monocular and binocular viewing for pose estimation in similar conditions. All experiments presented in this paper were conducted in compliance with the protocol approved by the Institutional Review Board at SUNY College of Optometry, and the Declaration of Helsinki, with observers giving written informed consent.

### Results

Perceived 3D lengths, averaged over 6 observers, are plotted against 3D pose for 5 different tilts of the display, separately for each of the three physical lengths in Figure 2a (Individual results in Figure A2). Dashed lines indicate the physical length of the test stick. In the fronto-parallel view, there is greater underestimation of length for poses pointing towards or away from the observer, and the longer the physical length of the test stick, the greater the underestimation of length. Increasing the tilt of the screen seriously reduced estimates of fronto-parallel sizes in oblique views, while there was little effect on perceived sizes of poses pointing towards or away from the observer. The videos in Figure 3, where rigid L-shaped figures are rotated in pose in frontal and oblique views, illustrate these two effects. The magnitudes of these two effects are easily seen in Figure 2b, where the fronto-parallel size estimate is subtracted from the oblique estimate, and the difference divided by the physical length. There are roughly equal magnitudes of underestimation as a fraction of the physical length for the three lengths, and similar patterns of underestimation as a function of object pose: the least change in estimates of 3D sizes caused by the tilt of the monitor are for poses close to the line of sight, whereas sizes of objects in poses close to fronto-parallel were seriously underestimated in oblique views compared to the frontal view. The effect of screen tilt also shows up in the pattern of underestimation for positive tilts being almost mirror symmetric to the pattern for negative tilts.

**Figure 2.**
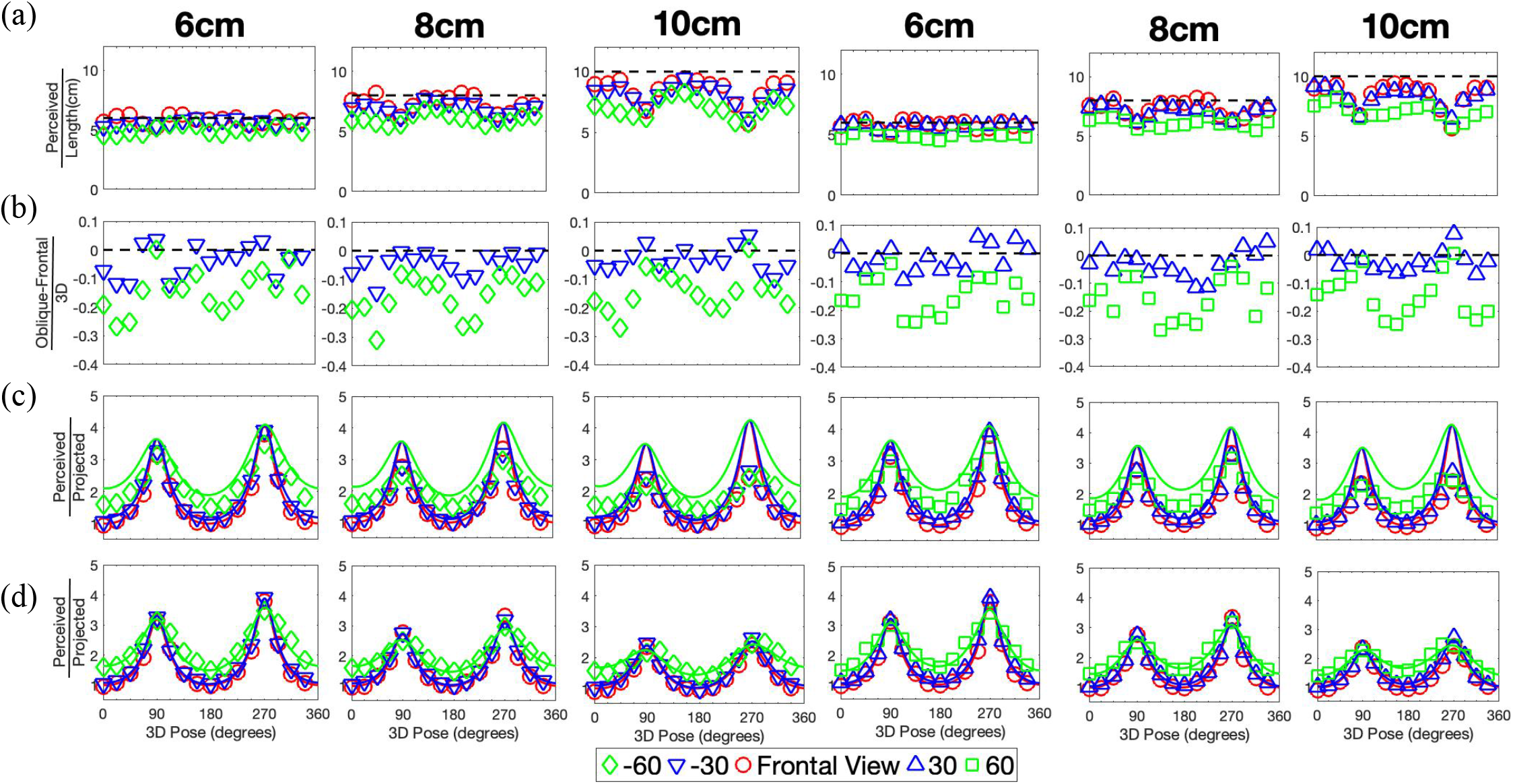
Average perceived 3D lengths (6 observers) Each column represents estimated lengths of a parallelepiped of the indicated physical length. Left three panels are for negative oblique viewing azimuths and right for positive. (a) Perceived length across 3D pose. Dashed lines indicate physical 3D length. Underestimation of perceived length increases systematically with increased tilt of the display. (b) Perceived 3D length for oblique view minus perceived length for frontal view divided by 3D physical length. The dotted line indicates no difference in perceived length from frontal view. (c) Optimal Correction Factor (solid line) and Empirical Correction Factor (symbols) across 3D pose, color coded for display tilt. Higher optimal correction is required at the fronto-parallel poses as display tilt increases, but is fairly constant for poses around the line of sight. (d) A Model using the optimal geometrical back-transform, but incorporating underestimation of the tilt of the display, fits the correction of perceived length over the projected length.

Since horizontal tilts of the screen lead to the greatest length compression along the fronto-parallel axis (Figure 1), the question arises whether the decrease in perceived length is explained completely by the shorter projected length, or whether the visual system does compensate partially for this compression. On the dark ground, the retinal image of the object holds the only information available for size estimation. Koch, Baig and Zaidi (2018) showed that in perception of poses in oblique views of pictures, the same observer-centered back-transform is used as for real scenes, thus leading to pose estimates that are a rigid rotation of actual scene poses by an angle equal to the viewing azimuth. Maruya and Zaidi (In press) showed that perceiving 3D sizes in real scenes also uses the optimal geometric back-transform for sizes, but estimates are sub-optimal because of a slant illusion that makes longer objects appear more slanted, which leads to less correction than required. The results in Figure 2 show that observers don’t make veridical estimates of 3D size. It is still possible that they use the optimal geometrical back-projection from the retinal image, but misestimate one or more parameters that are part of the back-transform expression. The physical 3D length divided by the length projected on the retina *L*_3*D*_/*L_r_* gives the Optimal Length Correction index (OLC) for each pose in each viewpoint (plotted as color-coded solid curves in Figure 2c). The symbols in Figure 2c plot the perceived lengths from Figure 2a divided by projected lengths giving the Measured Length Corrections (MLC). OLCs at line of sight are the highest with the same values regardless of the tilts of the display. The largest values of MLC also correspond to poses pointing towards or away from the observer, and these values are similar across different tilts of the display, but they are lower than what is required for veridical estimates, especially for the longer lengths. In contrast, OLCs at fronto-parallel poses increase with the tilt of the display. Similarly, MLCs at these poses increase with the tilt of display, but not enough. Figure 2c enables us to reject the hypothesis that size perception in pictures is similar to pose perception in using the same back-transform as for real scenes, irrespective of the tilt of the monitor. If this hypothesis was correct, the frontal view OLC, which is also the OLC for real scenes, should fit the data with some multiplicative scaling of the red curve, but that cannot happen because OLC for 0°and 180° poses is anchored at 1.0 when display tilt is fixed at 0°, no matter what the values for other parameters such as camera elevation, focal length or distance, whereas the MLC for 0°and 180 is higher for the tilted displays showing increasing length correction with increasing tilt of the display. This increase is still substantially less than required by the OLC for the +/− 60° display tilt. The general form of Measured Length Correction as a function of 3D pose is similar to the Optimal Length Correction curve, suggesting that observers may be using the optimal back-transform, but with additional multiplicative factors leading to the sub-optimality.

### Model

The geometrical back-transform is obtained by inverting Equation 1 to get an expression for the estimated 3D length 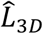:

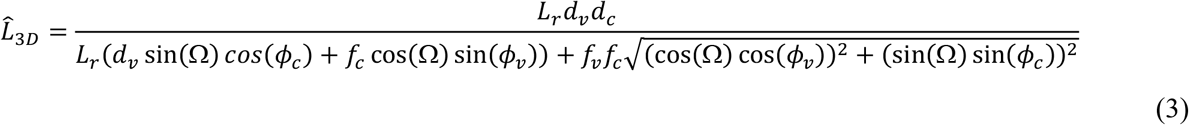

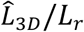 gives the expression for the Estimated Length Correction index (ELC) for each pose. The ELC will be equal to the OLC, and give veridical estimates of 3D size, if values of viewing elevation and azimuth are accurate. If these values are not accurate, the estimated 3D length 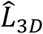 will be different from the veridical. We will try to understand the suboptimal size estimates in obliquely viewed pictures by seeing if they can be predicted by inaccurate estimates of display tilt, and then directly measuring perceived tilt. The motivation for this manipulation is the fronto-parallel bias in judging 3D pose (Koch, Baig Zaidi, 2018).

Maruya and Zaidi (In press) showed that perceived sizes can be inaccurate in 3D scenes despite observers using the correct geometric back-transform, if the retinal image evokes a misestimate of viewing elevation. The longer the physical length of the test stick, the greater the misestimation of the slant, and the effect is most obvious for poses pointing at or away from the observer. Their model fit the MLCs with just one free parameter that modified the estimated camera elevation (equivalent to modifying perceived slant). To illustrate the slant illusion, they made the video shown in the frontal view of Figure 4. When the arms of the figure from Figure 3 are dynamically adjusted to be perceptually equal in length across poses, observers perceive a change in slant every time an arm passes through the line of site. Figure 4 shows that the illusion persists when the display is tilted, so we retain this modification of the model. In addition, higher OLCs are required at fronto-parallel poses with increasing tilt of the display, but MLCs are uniformly lower than adequate. We thus include underestimation of the tilt of the display in the model, because ELCs are uniformly lower at the shallower tilts of the display.

**Figure 3.**
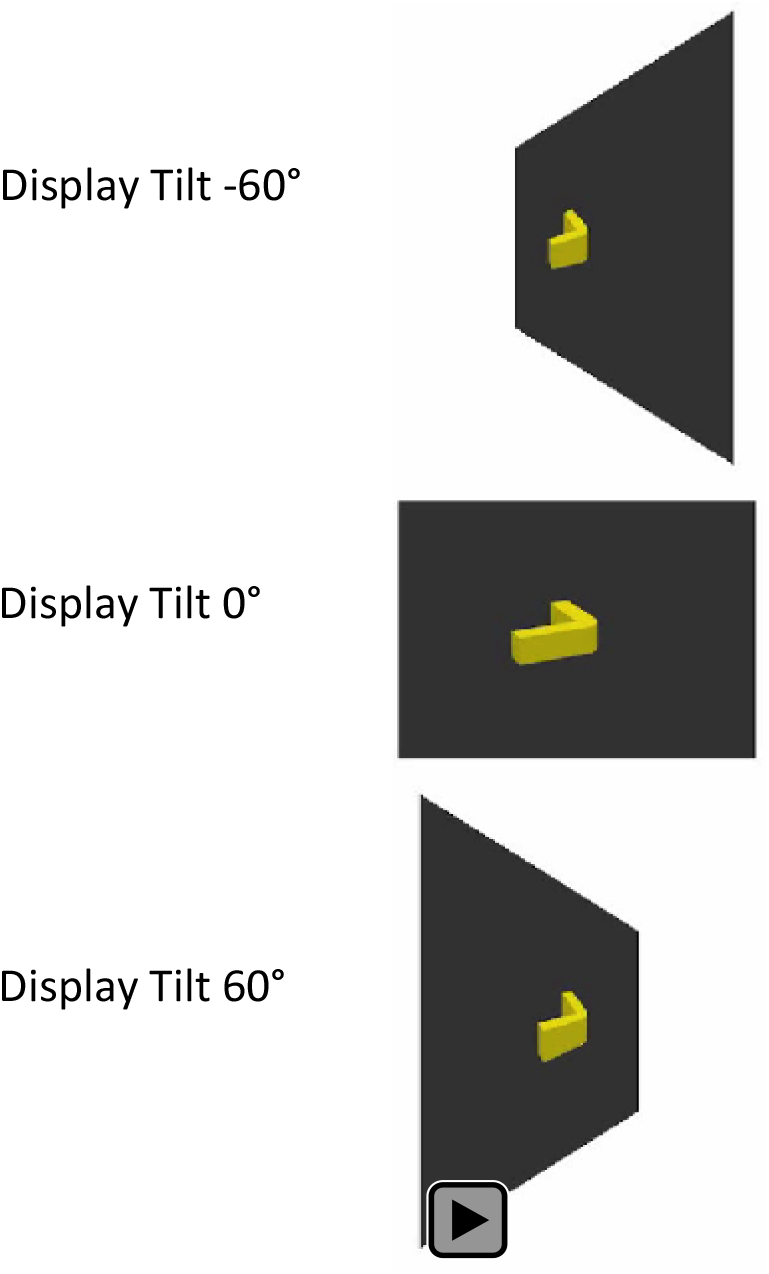
Dynamic demonstration of size inconstancy. A rigid object with two physically equal limbs at a right angle is rotating on the ground. When viewed from the front at a 15° elevation, the limb pointing at or away from the observer appears transitorily shorter than the orthogonal limb. Comparing the oblique views to the frontal view, the biggest change in length is transitorily for the fronto-parallel limb. The effects of the different signs of display tilt.

**Figure 4.**
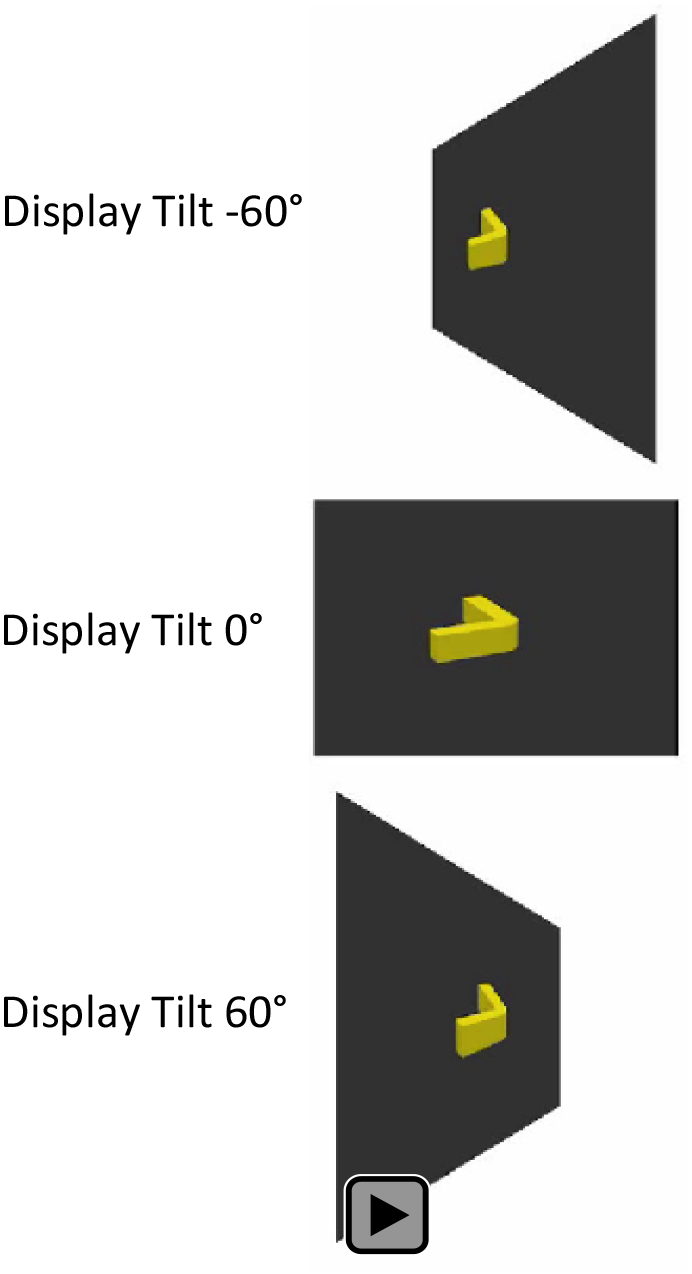
Dynamic demonstration of slant illusion. A rigid object with two limbs at a right angle is rotating on the ground. The length of the limb passing through the line of sight is lengthened and then shortened, to make the limbs appear equal in all poses in the view from the 15° elevation. The percept is maintained by increasing the length of the limb passing through the line of sight according to the average size estimate in Experiment 1. Instead of seeing the physical length changes, each limb seems to bounce up and down when it faces towards or away from the observer, because the slant illusion dominates the percept in the oblique views as well as the frontal view.

We formulated the hypothesis that observers are using an optimal back-transform, but overestimating the slant of the object (or equivalently the camera elevation) which leads to underestimated lengths at poses around 90° and 270°, plus they are underestimating the tilt of the display, thus correcting less than required for poses around 0° and 180° degrees. We tested whether adding multipliers *k_c_* > 1 to the viewing elevation, and *k_v_* < 1 to the tilt of the display, in the optimal geometrical back-transform expression, could provide good fits to the MLCs:

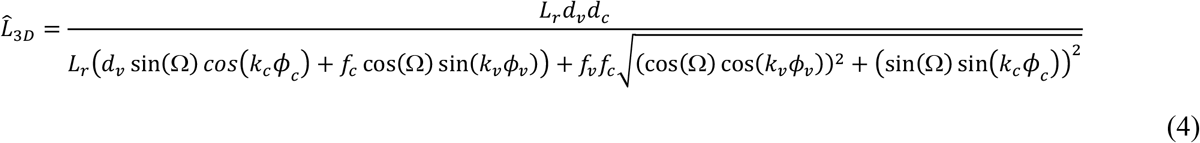

For each length and the tilt of the display, Figure 2(d) replots the empirical corrections from Figure 2(c) and predicted values of Equation 4 for the *k_c_* and *k_v_* that give the best least squares fit to the data. The model fits the results for all three lengths and tilts of the display well, with just the two free parameters (Best fits to individual results in Figure A3). Best fitting *k_c_* are nearly equal for fronto-parallel and oblique views, and best fitting *k_v_* correspond to perceived display tilts 21.3° for 30° view, −19.2° for −30°, 49.8° for 60°, −50.4° for −60°. Parenthetically, we explicitly tested whether simulating misperceived distance (Sedgwick, 1989) by putting multipliers on the distance, could explain the data, but that was not successful by itself or when combined with misperceived slant. To validate the predictions of the best fitting model, we measured directly whether observers actually misperceived the tilt of the display.

## Display Tilt Underestimation

### Methods

To test whether there was actual misestimation of the tilt of the display where the stimuli were presented, we presented a 10 cm test stimulus at 0° pose at the center of the display, and observers were instructed to judge the 3D pose of the screen. Viewing position was identical to Experiment 1, and the scene was viewed with tilts of 0°, ±30°, and ±60° in random order. Observers recorded their judgement by rotating a vector in a clock face on an iPad to the same angle as the pose of the rectangle. The iPad screen was placed close to the display screen, and observers adjusted the vector angle on a keyboard. Measurements were separated into three sets and observers were allowed to take a break between sets.

### Results

The main result (Figure 5) is that observers underestimated the tilt of the display at oblique viewing conditions despite binocular viewing. Average perceived tilts of the display were 23.24° for 30° view, −19.40° for −30°, 47.27° for 60°, and −49.47° for −60°. As we expected, the underestimation pattern for the obliquely viewed display surface is consistent with the fronto-parallel bias for obliquely posed objects (Koch et al, 2018). The results support our hypothesis that observers may be applying a smaller correction to oblique views because there is underestimation of the tilt of the display. Measured perceived tilts are similar in value to those predicted by the best fitting model.

**Figure 5.**
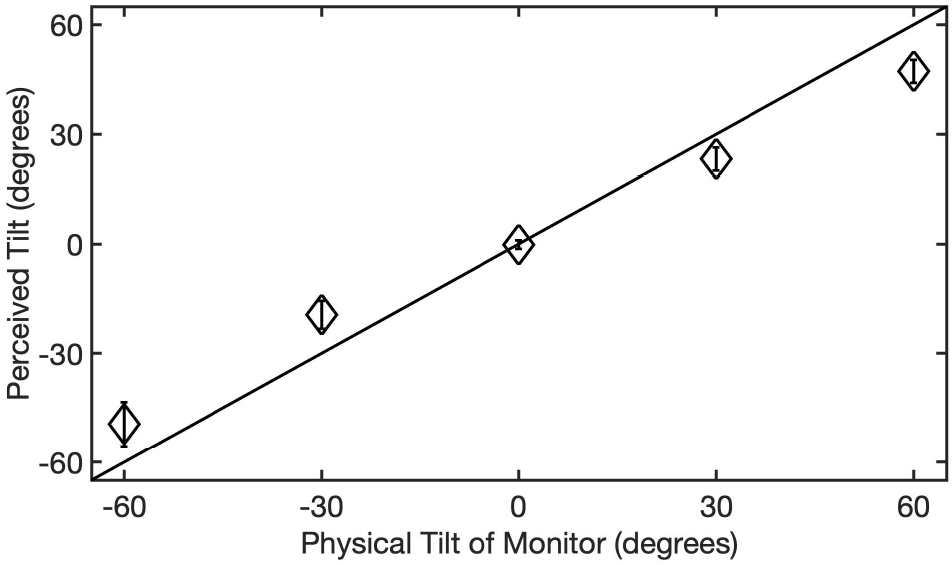
Display Tilt Underestimation. Perceived tilt as a function of physical tilt of monitor. In every oblique condition, there is underestimation of the tilt of the display.

## Discussion

Many authors have argued that picture perception is different from perception of real scenes, and pictorial space is different from real space (Kennedy, 1974; Ward, 1976; Pierroutsakos et al, 1998; Yang et al, 1999; Heyer, 2003; Vishwanath et al, 2005; Koenderink et al, 2004 and 2011; Pagel, 2017). Our approach to this issue is empirical and conceptual, as we examine visual tasks that are common to perceiving real and pictured scenes, and identify geometric operations that are involved in both perceptions. In judging poses of 3D objects, we found that observers judged poses in pictures by applying the same back-transform to retinal projections that they did for real scenes, and this predicted the illusory rotation of scenes to match the observer’s viewing azimuth. Correcting for the tilt of the picture would not predict the illusory rotation, so the case of object pose we see no reason to invoke special processes for picture perception. This study asks whether that is also true for judging 3D sizes in pictures. Although poses in pictures were invariant to viewing azimuth, there was noticeable distortion of lengths and aspect ratios. Display tilt reduces perceived sizes of frontal objects, and makes parallelepipeds pointing at the observer look narrower, which can be seen by comparing oblique to frontal views in Figures 1 and 4. Consequently, judgments of size and aspect ratio in pictures may be qualitatively different from judgments of poses, and may provide new considerations in picture perception.

The main empirical contribution of this paper is to measure the perceived size of 3D objects at different poses depicted on a planar picture rotated around its vertical axis. Pose variation and display tilt both shorten the projected length of 3D objects, but along orthogonal dimensions, pose variations along the line of sight, and display tilt along the fronto-parallel axis. Our results show that the visual system uses the geometric back-transform to overcome these distortions, but falls short because it overestimates slants for poses along the line of sight, and underestimates the tilt of the display. The same factors lead to perceived changes in aspect ratios of shapes, so that objects pointing towards the observer look narrower in oblique views of pictures.

The main theoretical contribution of this study is to link perception of 3D sizes in scenes and pictures to the mental use of projective geometry. Corrections of sizes from projective distortions as a function of pose form a curve that has the same shape as that predicted by the optimal back-transform, and the optimal correction expression fits measured estimates with just two free parameters multiplying the camera angle by greater than 1.0 and the display tilt by less than 1.0. Thus, our model that incorporates observers’ misestimates of object slant and display tilt can explain the inconstancy of relative size for different poses, object sizes, and viewpoint azimuths, suggesting that the mental use of projective geometry is common to all observers.

Animals and humans have constant exposure to perspective projection through image-forming eyes. Therefore, whether brains have learned to exploit projective geometry to understand real scenes is an age old question, e.g. Plato’s dialogue Meno. We have now shown that humans use optimal projective geometry back-transforms from retinal images to estimate 3D pose and size in real scenes, and continue to use the back-transforms to estimate 3D pose and size in pictures, despite the extra distortions created by oblique views of pictures (Koch, Baig & Zaidi, 2018; Maruya & Zaidi, In press). These results provide accumulating evidence that human brains have internalized particular aspects of projective geometry through evolution or learning.

## ACKNOWLEDGMENTS

This work was supported by National Institutes of Health Grants EY13312 and EY07556.

## AUTHOR CONTRIBUTIONS

AM and QZ designed the study. AM programmed and ran the experiments. AM & QZ analyzed and modeled the results. AM & QZ wrote the paper.

The authors declare no competing interests

### Precis

Observers estimate 3D sizes in oblique views of pictures by using internalized projective geometry, but correct inadequately for projected shortening because of underestimating picture tilt.

## Appendix

## Derivation of 2D retinal lengths from 3D lengths of object at poses

This projection is derived for the blue parallelepiped lying on the ground plane in 3-Space (UVW space, where the X-Z face is on the top surface of the parallelepiped and the Y-axis is the center of the rotation), and extending from the center of the scene (0,0,0) to (*x*, 0, *z*). The end-point (*x*,0,*z*) can be expressed in terms of the physical length (*L*_3*D*_) and pose (Ω) of the parallelepiped:

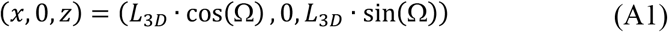

Since the camera elevation, *ϕ_c_*, from the Z-axis is equivalent to a rotation around the X-axis, the rotation of the end-point (*x*, 0, *z*) is given by:

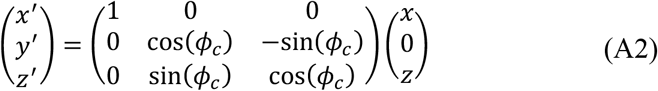

So that:

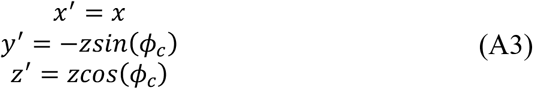

The center point stays the same, (0,0,0) → (0,0,0).

In the projection to the Picture Plane (UV-Space), the central point is mapped as (0,0,0) → (0,0), and the coordinates of the end-point are:

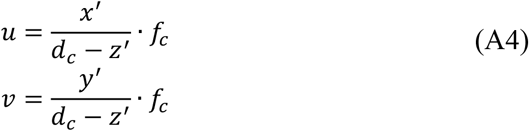

So the projected length on the Picture Plane is given by:

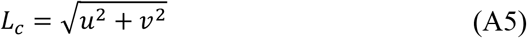

By substituting Equation (1), (3), and (4) into (5)

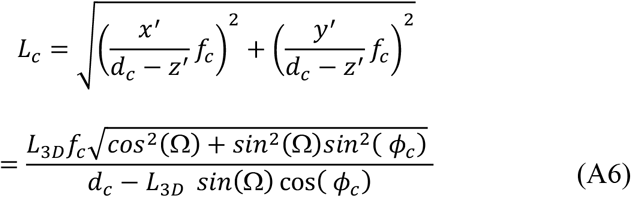

Finally the Picture Plane is projected to the retinal plane (RS-space). Since the Picture Plane is tilted by *ϕ_v_* (±30° *or* ± 60°), while the observer’s viewing position is fixed, the new coordinates are defined in 3-Space (UVW space, where the W axis is orthogonal to the fronto-parallel location of the screen, adding depth to the 2D Space defined by the fronto-parallel picture plane. The central point is mapped as (0,0) → (0,0,0). The new coordinates for the end-point are:

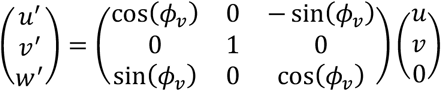

Giving:

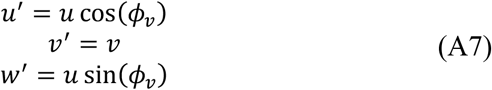

If *d_v_* is the distance from the observer’s pupil to the center of the picture plane, and *f_v_* is the observer’s focal length, the end-point’s projection from UVW-Space to retinal space is:

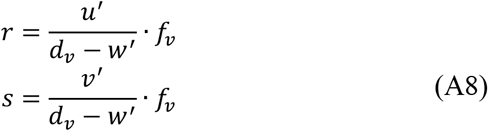

Therefore, the projected length on the retina:

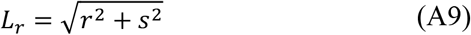

is derived by substituting Equation (1), (3), (4), (7), and (8) into (9)

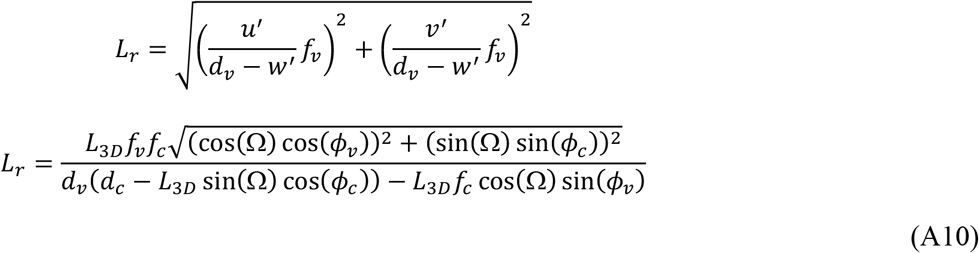

## Figure legends (Appendix)

**Figure A1:**
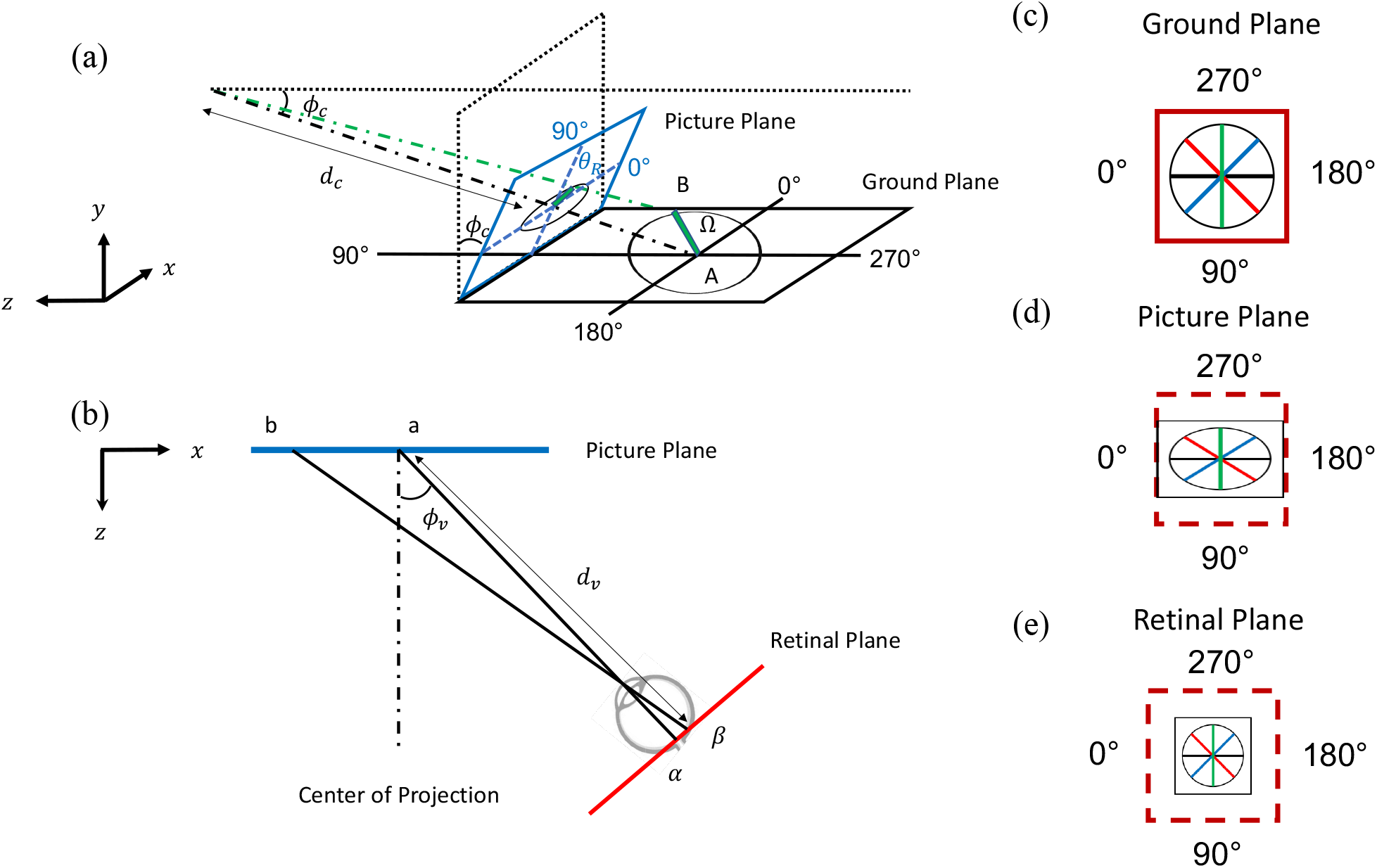
Derivation of projected length as a function of object pose and viewing azimuth. (a) Equal lengths of objects on the ground project to different lengths on the screen as a function of pose, as a circle is compressed into an ellipse. Comparing panel (c) to (d) shows greatest reduction for line of sight poses. (b) When the screen is viewed obliquely, in the retinal projection, the ellipse is compressed towards a circle again. Comparing panel (d) to (e) shows the reduction in length is greatest for fronto-parallel poses.

**Figure A2.**
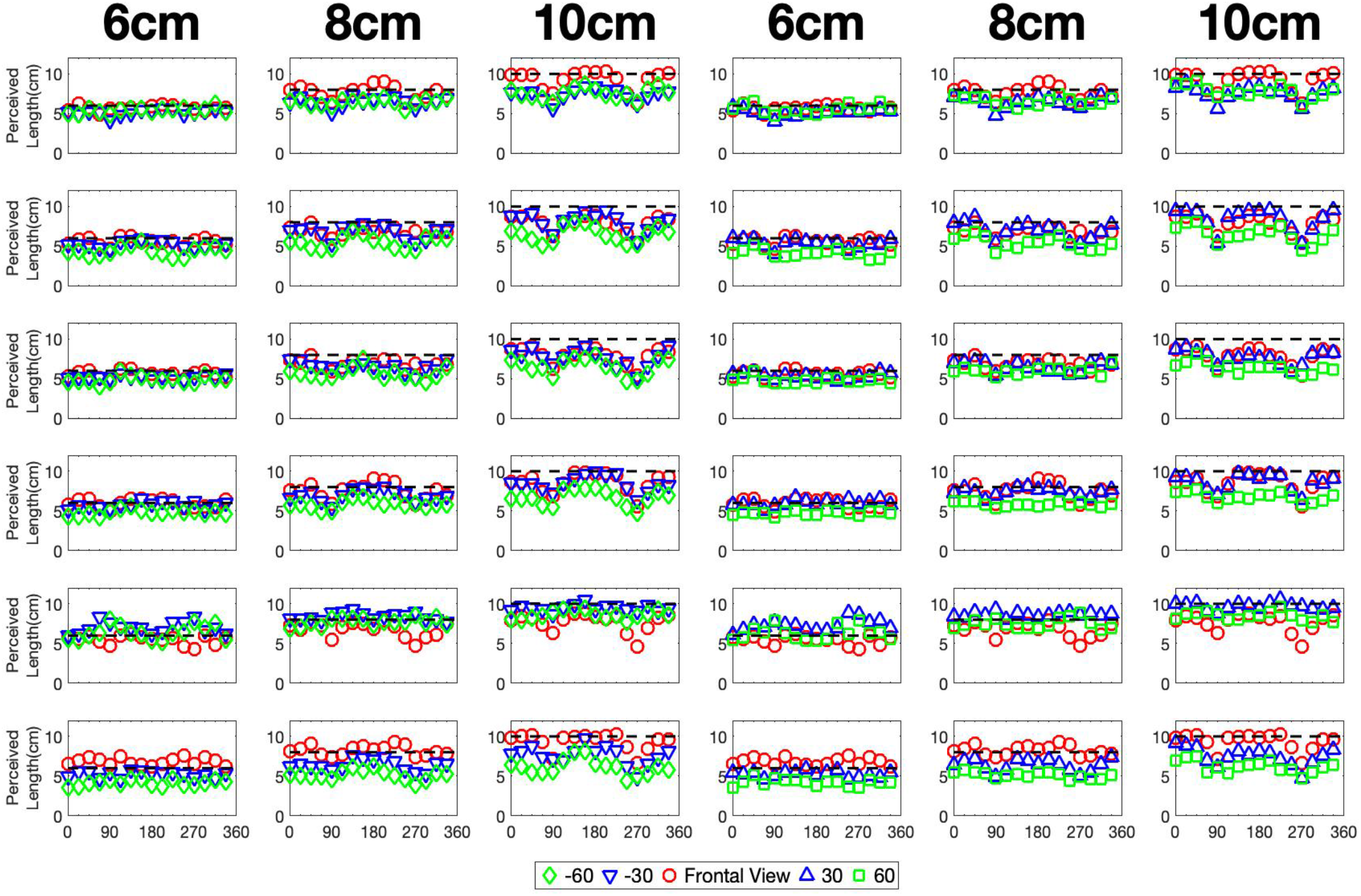
Perceived 3D lengths across pose and display slant for 6 observers. Each column represents estimated lengths of a parallelepiped of the indicated physical length. Left three panels are for negative oblique viewing azimuths and right for positive. Points show perceived length across 3D pose. Dashed lines indicate physical 3D length. Underestimation of perceived length increases systematically with increased tilt of the display.

**Figure A3.**
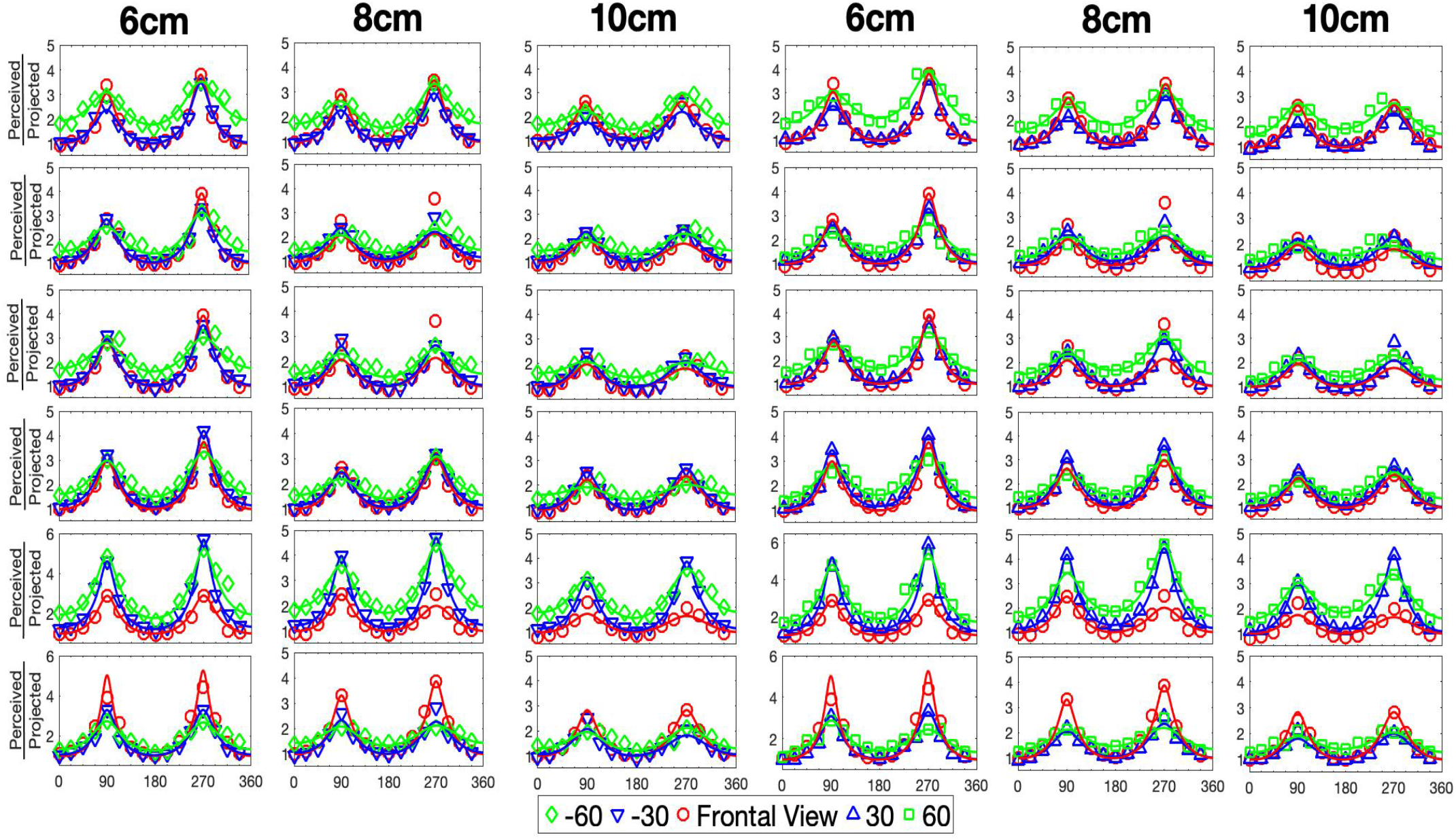
Model fits for 3D length estimation across pose and display slant for 6 observers. Each column represents estimated lengths of a parallelepiped of the indicated physical length. Left three panels are for negative oblique viewing azimuths and right for positive. A Model using the optimal geometrical back-transform, but incorporating underestimation of the tilt of the display, fits the correction of perceived length over the projected length.

**Figure A4.**
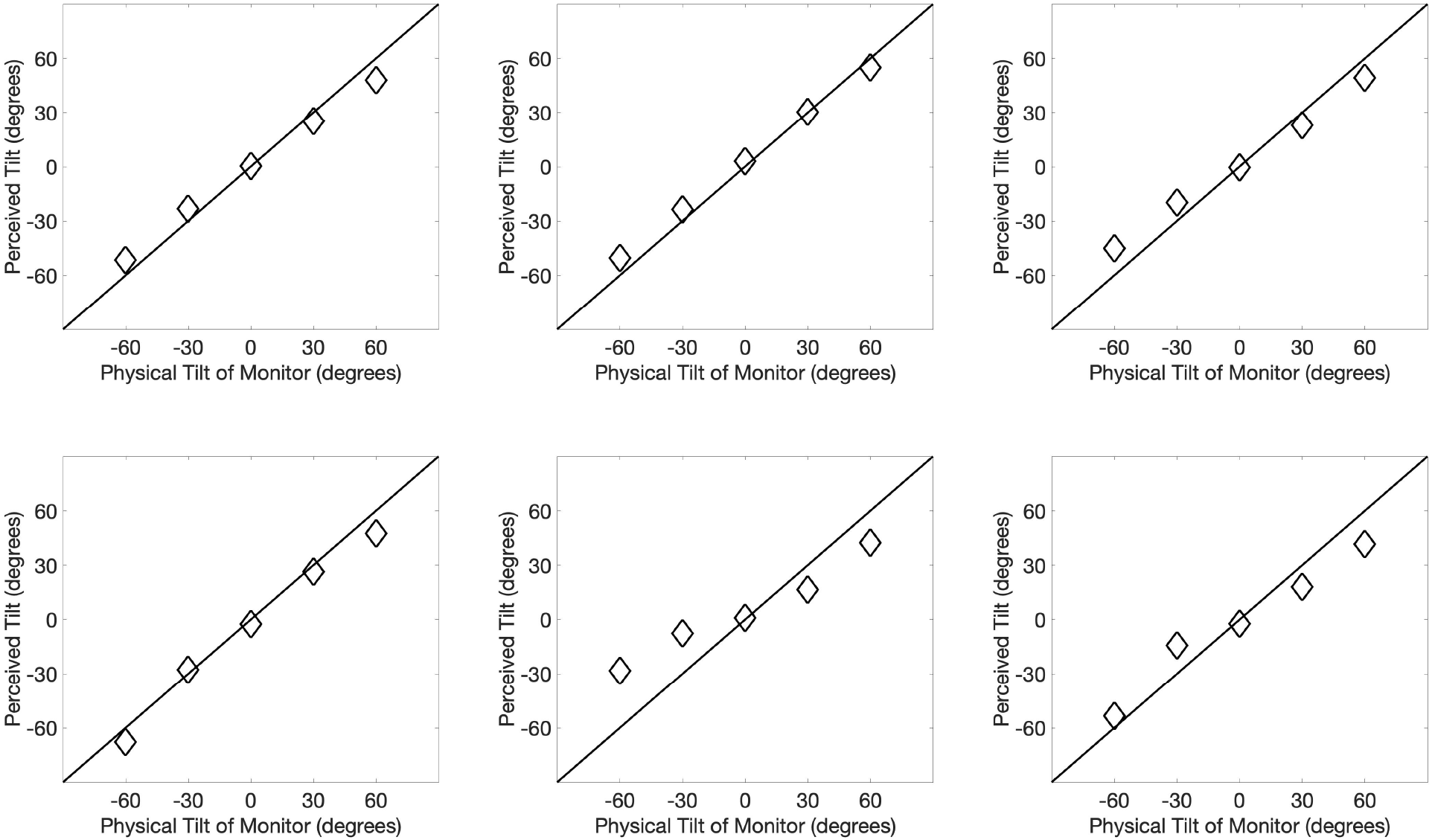
Perceived Display Tilt for 6 observers. Perceived display tilt as a function of physical tilt of monitor for 6 observers. In almost every oblique viewing condition, there is underestimation of the tilt of the display.

## References

Beusmans, J. M. (1998). Optic flow and the metric of the visual ground plane. Vision research, 38(8), 1153–1170.

Boring, E. G. (1964). Size-constancy in a picture. The American journal of psychology, 77(3), 494–498.

Brunswik, E. (1944). Distal focussing of perception: Size-constancy in a representative sample of situations. Psychological Monographs, 56(1), i.

Carlson, V. (1960). Overestimation in size-constancy judgments. The American journal of psychology, 73(2), 199–213.

Cutting, J. E. (1987). Rigidity in cinema seen from the front row, side aisle. Journal of Experimental Psychology: Human Perception and Performance, 13(3), 323.

DeLoache, J. S., Pierroutsakos, S. L., Uttal, D. H., Rosengren, K. S., & Gottlieb, A. (1998). Grasping the nature of pictures. Psychological Science, 9(3), 205–210.

Gilinsky, A. S. (1951). Perceived size and distance in visual space. Psychological review, 58(6), 460.

Gilinsky, A. S. (1955). The effect of attitude upon the perception of size. The American journal of psychology, 68(2), 173–192.

Gombrich, E. H. (1972). The visual image. Scientific American, 227(3), 82–97.

Hagen, M. A. (1976). Influence of picture surface and station point on the ability to compensate for oblique view in pictorial perception. Developmental Psychology, 12(1), 57.

Kennedy, J. M. (1974). A psychology of picture perception: Jossey-Bass Publishers.

Koch, E., Baig, F., & Zaidi, Q. (2018). Picture perception reveals mental geometry of 3D scene inferences. Proceedings of the National Academy of Sciences, 115(30), 7807–7812.

Koenderink, J. J., van Doom, A. J., Kappers, A. M., & Todd, J. T. (2004). Pointing out of the picture. Perception, 33(5), 513–530.

Koenderink, J. J., van Doorn, A. J., & Wagemans, J. (2011). Depth. i-Perception, 2(6), 541–564.

Loomis, J. M., Da Silva, J. A., Fujita, N., & Fukusima, S. S. (1992). Visual space perception and visually directed action. Journal of Experimental Psychology: Human Perception and Performance, 18(4), 906.

Loomis, J. M., & Philbeck, J. W. (1999). Is the anisotropy of perceived 3-D shape invariant across scale? Perception & Psychophysics, 61(3), 397–402.

Loomis, J. M., Philbeck, J. W., & Zahorik, P. (2002). Dissociation between location and shape in visual space. Journal of Experimental Psychology: Human Perception and Performance, 28(5), 1202.

Maruya, A., & Zaidi, Q. (2020). “Mental geometry of 3D size perception.” Journal of vision (In press)

Niall, K. K., & Macnamara, J. (1990). Projective invariance and picture perception. Perception, 19(5), 637–660.

Niederée, R., & Heyer, D. (2003). The dual nature of picture perception: A challenge to current general accounts of visual perception. Looking into pictures: An interdisciplinary approach to pictorial space, 77–98.

Norman, J. F., Todd, J. T., Perotti, V. J., & Tittle, J. S. (1996). The visual perception of threedimensional length. Journal of Experimental Psychology: Human Perception and Performance, 22(1), 173.

Pagel, R. (2017). The duality of picture perception and the robustness of perspective. Art & Perception, 5(3), 233–261.

Perkins, D. N. (1973). Compensating for distortion in viewing pictures obliquely. Perception & Psychophysics, 14(1), 13–18.

Rosinski, R. R., Mulholland, T., Degelman, D., & Farber, J. (1980). Picture perception: An analysis of visual compensation. Perception & Psychophysics, 28(6), 521–526.

Ross, H. E., & Plug, C. (1998). The history of size constancy and size illusions.

Sedgwick H. A. (1989). “The effects of viewpoint on the virtual space of pictures.” Available at https://ntrs.nasa.gov/search.jsp?R=19900013616.

Todorović, D. (2008). Is pictorial perception robust? The effect of the observer vantage point on the perceived depth structure of linear-perspective images. Perception, 37(1), 106–125.

van Doorn, A. J., Koenderink, J. J., Leyssen, M. H., & Wagemans, J. (2012). Interaction of depth probes and style of depiction. i-Perception, 3(8), 528–540.

Vishwanath, D., Girshick, A. R., & Banks, M. S. (2005). Why pictures look right when viewed from the wrong place. Nature neuroscience, 8(10), 1401–1410.

Wallach, H., & Marshall, F. J. (1986). Shape constancy in pictorial representation. Perception & Psychophysics, 39(4), 233–235.

Ward, J. L. (1976). The perception of pictorial space in perspective pictures. Leonardo, 9(4), 279–288.

Yang, T., & Kubovy, M. (1999). Weakening the robustness of perspective: Evidence for a modified theory of compensation in picture perception. Perception & Psychophysics, 61(3), 456–467.

